# Application of Pharmacogenomics and Bioinformatics to Exemplify the Utility of Human *ex vivo* Organoculture Models in the Field of Precision Medicine

**DOI:** 10.1101/649376

**Authors:** K Cowan, G Macluskie, M Finch, C.N.A Palmer, J Hair, M Bylesjo, S Lynagh, P Brankin, M McNeil, C Low, D Mallinson, EM Gourlay, H Child, L Cheyne, DC Bunton

## Abstract

Here we describe a collaboration between industry, the National Health Service (NHS) and academia that sought to demonstrate how early understanding of both pharmacology and genomics can improve strategies for the development of precision medicines. Diseased tissue ethically acquired from patients suffering from chronic obstructive pulmonary disease (COPD), was used to investigate inter-patient variability in drug efficacy using *ex vivo* organocultures of fresh lung tissue as the test system. The reduction in inflammatory cytokines in the presence of various test drugs was used as the measure of drug efficacy and the individual patient responses were then matched against genotype and microRNA profiles in an attempt to identify unique predictors of drug responsiveness. Our findings suggest that genetic variation in CYP2E1 and SMAD3 genes may partly explain the observed variation in drug response.

## Introduction

It is well recognised that one size does not fit all when it comes to the treatment of many diseases. Getting the right drug to the right patient at the right dose has become the focus of precision medicine, which provides hope that patients may receive the most appropriate treatment sooner, improving their quality of life and reducing the support required from health care systems and wider society^1^. Health economists are recognising the potential of precision medicine and are beginning to apply the concept to their research.^2^

The genomics revolution has underpinned much of this research. As the cost of gene sequencing has fallen, the ability to rapidly identify an individual’s genotype as part of routine health care has become possible. However, for precision medicines to be developed, genomics must be linked to pharmacology: relating the individuals genotype to the effectiveness, potency and tolerability of a drug. It is through pharmacogenomics that truly personalised therapies may emerge, yet the link between genomics and pharmacology may not be properly understood until expensive and risky clinical trials are conducted.

Here we describe a collaboration between industry, the National Health Service (NHS) and academia that sought to demonstrate how early understanding of both pharmacology and genomics can improve strategies for the development of precision medicines. By using the latest pharmacology techniques in human fresh tissues from the target patient population, combined with genomics and clinical metadata associated with each individual, an improved understanding of the link between genetics and inter-individual drug responses emerges.

An early understanding of patient stratification during drug discovery is becoming increasingly important. Selection and optimisation of candidate drugs for well-defined patient subsets has the potential to help in the design of more rapid, targeted clinical trials.

A key incentive to better understand pharmacogenomics during the drug discovery process is the rapid increase in drug development costs. The most recent estimates of the out-of-pocket costs (i.e. excluding capital costs) of drug development are in the region of $890m^7^, with approximately 70% of the costs incurred during clinical development. The most common cause of failure is poor efficacy at phase II or III^3.4.5.6^, which is in part attributed to trials of entire patient populations that include both “responders” and “non-responders”. Precision medicine can improve the prediction of clinical efficacy by selecting for clinical trials only those patient sub-populations likely to gain clear benefit; such predictions are dependent on the quality of the information used to stratify the patient sub-populations at an early stage of development. Early data on the effectiveness of drugs in different patients is essential to the development of precision medicines. Pre-clinical tests of drug effects must therefore closely reflect the patient population.

The most desired traits in pre-clinical models are “physiological relevance” and the ability to translate findings to likely clinical responses^3, 6–8^, including a desire to model the likely variation in effectiveness of a new drug within the patient population. Human fresh tissues and complex 3D tissue models that reflect the biology of disease are therefore increasingly being used by Pharma to improve the prediction of efficacy in clinical trials^7, 9, 10^. Although the data between different patients can be variable, this is now viewed as an opportunity for an early understanding of the extent and causes of inter-patient variation in drug response.

Chronic Obstructive Pulmonary Disease (COPD) is a major health problem and is an example of a complex condition, with many clinical phenotypes. Many patients receive minimal clinical benefit from common medications, most likely due to the combination of variations in disease subtype and genotype. The clinical variation in drug response is apparent in *ex vivo* pharmacology experiments using fresh lung tissues^29^.

In this project, diseased tissue ethically acquired from patients suffering from COPD, was used to investigate inter-patient variability in drug efficacy using *ex vivo* organocultures of fresh lung tissue as the test system. In order to assess patient variation and responsiveness to both ‘standard of care’ and potential novel therapies, the reduction in inflammatory cytokines in the presence of various test drugs was used as the measure of drug efficacy. The individual patient responses were then matched against genotype and microRNA profiles in an attempt to identify unique predictors of drug responsiveness and demonstrate the combined power of pharmacology and genomics during pre-clinical development.

## Materials and methods

### Organoculture - REPROCELL

#### Materials

RPMI 1640 glutamax culture medium, gentamicin (50 mg/ml) and amphotericin B (250 µg/ml) were purchased from Thermo Fisher Scientific. Retinyl acetate, nystatin, bovine insulin, foetal bovine serum (FBS), fluticasone, roflumilast, RNAlater and DMSO were purchased from Sigma. Formoterol was purchased from R&D Systems and lipopolysaccharide endotoxin (LPS) was purchased from Invivogen. Complete mini protease inhibitor was purchased from Roche.

#### Method

COPD lung parenchyma tissue was ethically obtained from 25 patients undergoing therapeutic resection for cancer or COPD. Residual tissue, not required for diagnosis, was acquired from NHS Research Scotland Bio-repository Network and also through the REPROCELL tissue network. Patients provided written consent, complying with the declaration of Helsinki.

Lung parenchyma was dissected free from pleura, visible airways and blood vessels to produce 5 mm^3^ biopsies. Two biopsies were immediately placed in RNAlater and stored at 2 to 8°C overnight, prior to storage at –80°C. Remaining biopsies were subjected to the following culture protocol.

RPMI 1640 culture medium was prepared by adding the following constituents: gentamicin (100 µg/ml), amphotericin B (0.625 µg/ml), FBS (0.5%), retinyl acetate (0.1 µg/ml), bovine insulin (1 µg/ml) and nystatin (1 µg/ml). Final concentration of each constituent is displayed.

Biopsies were submerged in culture media (two biopsies per well) and incubated for 16 to 24h at 37 °C, in the presence of 5% CO_2_.

Following the incubation period, media was refreshed and each well containing two biopsies was exposed to LPS (100 ng/ml), in an attempt to boost and normalise inter-biopsy inflammatory cytokine release. Each well was assigned one of the following experimental conditions: DMSO vehicle; roflumilast (100 nM); fluticasone (1 µM); formoterol (10 nM); or a combination of roflumilast (100 nM) plus fluticasone (1 µM) or formoterol (10 nM). Biopsies were then subjected to a further incubation period of 24 h at 37 °C in the presence of 5% CO_2_. Culture supernatants were sampled from each well, protease inhibitor added to prevent degradation of inflammatory cytokines and stored at −80°C prior to analysis.

Each experimental condition was performed in duplicate culture wells.

#### Luminex MAGPIX Analysis

Levels of TNFα (pg/ml) were measured in culture supernatants using a magnetic bead-based assay for the Luminex MAGPIX platform. Fluorescence levels correlating with TNFα level were corrected against the blank control level and a standard curve was generated using a 5-parameter logistic equation.

Each culture supernatant sample was analysed in duplicate and reported by Bio plex Manager 6.1 software as mean TNFα concentration (pg/ml), along with standard deviation of the mean and the percentage coefficient of variation.

Graph Pad Prism 4 software was used to display the data for all 25 donors as a median or mean TNFα concentration (pg/ml) + Standard Error of Mean (SEM.). Median and mean TNFα concentration was also displayed in Graph Pad Prism 4 as a percentage of DMSO vehicle control.

### RNA, DNA extraction and miRNA analysis – Sistemic

Two baseline lung biopsies were prepared from 25 donors, as described above, and transported to Sistemic for DNA and RNA extraction.

DNA was extracted from approximately 10 mg tissue using the PureLink Genomic DNA Mini Kit (Life Technologies). DNA quality control was performed using the Agilent 2200 TapeStation and the Genomic DNA ScreenTape kit to determine the DNA integrity number (DIN).

RNA was extracted from approximately 10 mg tissue. Tissue was homogenised in lysis buffer using a Precellys 24 homogeniser (Bertin Technologies) and total RNA was then extracted using the miRCURY RNA Isolation Kit – Cell & Plant (Exiqon). Absorbance ratios at 260/280 nM and 260/230 nM were determined as indicators of sample yield and purity.

Further RNA quality control was performed using the Agilent 2200 TapeStation and the ScreenTape R6K kit to determine the RNA integrity number (RIN).

MicroRNA (miRNA) expression levels were measured using the Agilent miRNA platform, specifically; Agilent’s SurePrint G3 Human v16 microRNA 8×60K microarray slides, miRBase version 16.0. Each slide contained 8 individual arrays and each array represents 1,349 microRNA’s; 1205 human miRNAs (mapped to 1194 miRNAs in miRBase 20) and 144 viral miRNAs.

Data was normalised using the AgiMicroRNA package in Bioconductor^11^. Array quality control was performed using outlier testing based on the following: average signal per array; average background per array; percentage of miRNAs where expression is detected on each array and the data distribution of each sample.

A sample to sample correlation analysis was performed on normalised data using Pearson’s correlation. Outliers were assessed using Grubbs’ outlier test with a significance threshold of p <0.05^12^.

miRNA expression data was visualised by Principal Component Analysis^13^, Pearson correlation and by agglomerative clustering heat-map in Bioconductor^14^. Isolated DNA was transported from Sistemic to the Stratified Medicine Scotland Innovation Centre (SMS-IC).

### Exome Sequencing – SMS-IC

Targeted next generation sequencing libraries were prepared using the Ion Ampliseq^TM^ Exome RDY Kit and DNA isolated from baseline lung biopsies. 60,496,505 bases were targeted by 293,903 amplicons, representing the coding sequence of 18,835 genes. Multiplexed PCR was performed to produce barcoded libraries, using 100ng of input DNA per sample and 10 amplification cycles. The Ion AmpliSeq^TM^ Library Kit Plus and IonXpress^TM^ Barcode Adapters were used in library preparation, according to the manufacturer’s instructions. Final library concentrations were determined by quantitative real time PCR using the Ion Library TaqMan^TM^ Quantitation Kit. Libraries were diluted to 100pM, and 2 libraries were subsequently pooled in equal amounts for templating on the Ion OneTouch^TM^ 2 System, using the Ion PI^TM^ Hi-Q^TM^ OT2 200 kit. The Ion Proton^TM^ NGS platform was used for sequencing of multiplexed templated libraries, using the Ion PI^TM^ Hi-Q^TM^ Sequencing 200 Kit and the Ion PI^TM^ Chip Kit v3, according to the manufacturer’s instructions.

### Raw Data Storage and Analysis – Aridhia & Fios Genomics

Raw data (organoculture TNFα response levels, miRNA expression profiles and exome sequencing data) was uploaded to a secure workspace (AnalytiXagility) in Aridhia’s digital research platform.

Anonymised, patient demographic data, obtained from NHS Research Scotland Bio-repository Network or the REPROCELL tissue network was also uploaded to the collaboration’s AnalytiXagility workspace. Data could then be accessed and analysed in a secure manner by authorised users.

Fios Genomics accessed data held in the AnalytiXagility research workspace to provide bioinformatic analyses. Each dataset was analysed individually and combined to determine any significant correlations between patient demographic data, genetic polymorphisms and/or miRNA profiles and the observed organoculture assay response.

#### Organoculture bioinformatic analysis

TNFα levels determined for each patient sample in the organoculture assay, were subjected to quality control metrics from the ArrayQualityMetrics package in Bioconductor^15^. Assays were scored on the basis of the following parameters: maplot; boxplot and heatmap. An individual patient sample was classified as an outlier if two or more of the parameters were not met.

TNFα levels (pg/ml) for each experimental condition were normalised using log_2_ ratios against DMSO vehicle control. Relative levels of TNFα were then visualised using bar charts, density plots and correlation plots within R software. The aim was to identify subgroups of patients that displayed a good reduction of TNFα levels in response to one or more of the organoculture experimental conditions and subgroups of patients that displayed a poor reduction.

Patients were then categorised as being a high responder or a low responder for use in subsequent bioinformatics analyses.

Defined patient demographic parameters and organoculture assay response were assessed using pair-wise univariate associations between all combinations of defined parameters. Associations between categorical parameters were assessed using a chi-squared test; associations between one categorical and one continuous parameter were assessed using analysis of variance (ANOVA); associations between two continuous parameters were assessed using a Spearman correlation test.

#### Exome sequence bioinformatic analysis

Torrent Mapping Alignment Program was used to provide IonTorrent AmpliSeq exome sequencing data for each patient. Data was provided as a BAM file aligned to genome reference GRCh37. Genotypes called with Torrent Variant Caller were provided as per sample VCF files.

Single nucleotide polymorphisms (SNPs) from the VCF files were merged into a multi-sample VCF and BAM files were used to set missing genotypes to homozygous reference if the read-depth of the SNP in a particular sample was less than 30. VCF files were then filtered to remove low quality SNPs.

Exploratory analysis was first performed by producing principal component analysis plots, using the SNPRrelate R software package. Hierarchical clustering of the data measured dissimilarity between patient exome data^16^.

The genotype for all SNPs identified from the VCF file was tested for association with the organoculture assay response, this was performed using fisher-exact tests of association within the Plink analysis toolkit^31^. Identified SNPs included those that were known to be related to genes of interest and also novel, undescribed SNPs.

Genes of interest were identified due to a literature association with the pathology of COPD and/or as being associated with lung metabolism and/or genes that may be associated with clinical response to standard of care treatments.

Identified SNPs were also cross-referenced with SNPs listed in the Genome Wide Association Studies (GWAS) Catalog to determine if any SNP had been previously reported in a human GWAS study and, if so, it was determined if the reported association was relevant to this study.

#### miRNA bioinformatic analysis

Quality control was assessed using the quality control metrics from the ArrayQualityMetrics package in Bioconductor^15^ as for the organoculture assay data above.

Confounding associations between defined patient demographic parameters and miRNA expression array data were assessed using pair-wise univariate associations between all combinations of defined parameters. Associations between categorical parameters were assessed using a chi-squared test; associations between one categorical and one continuous parameter were assessed using analysis of variance (ANOVA); associations between two continuous parameters were assessed using a Spearman correlation test.

Data was normalised using quantile normalisation which produces expression measures in a log base 2 format. Array batch effects due to processing of microarray data in two separate batches were corrected using the ComBat method^30^.

Statistical comparisons were performed to determine if specific miRNAs were associated with organoculture assay response: the null hypothesis being that no specific differences in miRNA expression could be detected in patients that responded well in the organoculture assay compared with patients that did not respond. Linear modelling, empirical Bayesian analysis and p-value adjustment for multiple testing (Benjamini-Hochberg) was performed using the Bioconductor Limma software package^14^.

miRNAs were annotated based on their experimentally verified target genes from miRTarBase^32^. miRNAs that displayed significant differential expression (uncorrected p <0.05), were analysed for enrichment of target gene KEGG (Kyoto Encyclopaedia of Genes and Genomes) pathway membership using a hypergeometric test. Upregulation and downregulation of genes were analysed separately.

In the same way, miRNA target genes were analysed for enrichment of gene ontology terms.

Integration of patient demographic, TNFα organoculture response, exome sequence and miRNA expression data

Congruence analysis was performed by evaluating the level of overlap between all data sets. Calculations of significant overlaps were based on a hypergeometric test.

All bioinformatic analysis was reviewed by Professor Colin Palmer, University of Dundee.

## Results

### Organoculture Luminex

The majority of COPD patient lung samples responded to treatment with fluticasone, roflumilast or combination therapy. This was observed as a reduction in the level of TNFα released from the biopsies into the culture media (Fig 2). Different levels of response were however observed between patients and ranged from modest to a marked reduction in TNFα in the supernatant.

**Figure 1.**
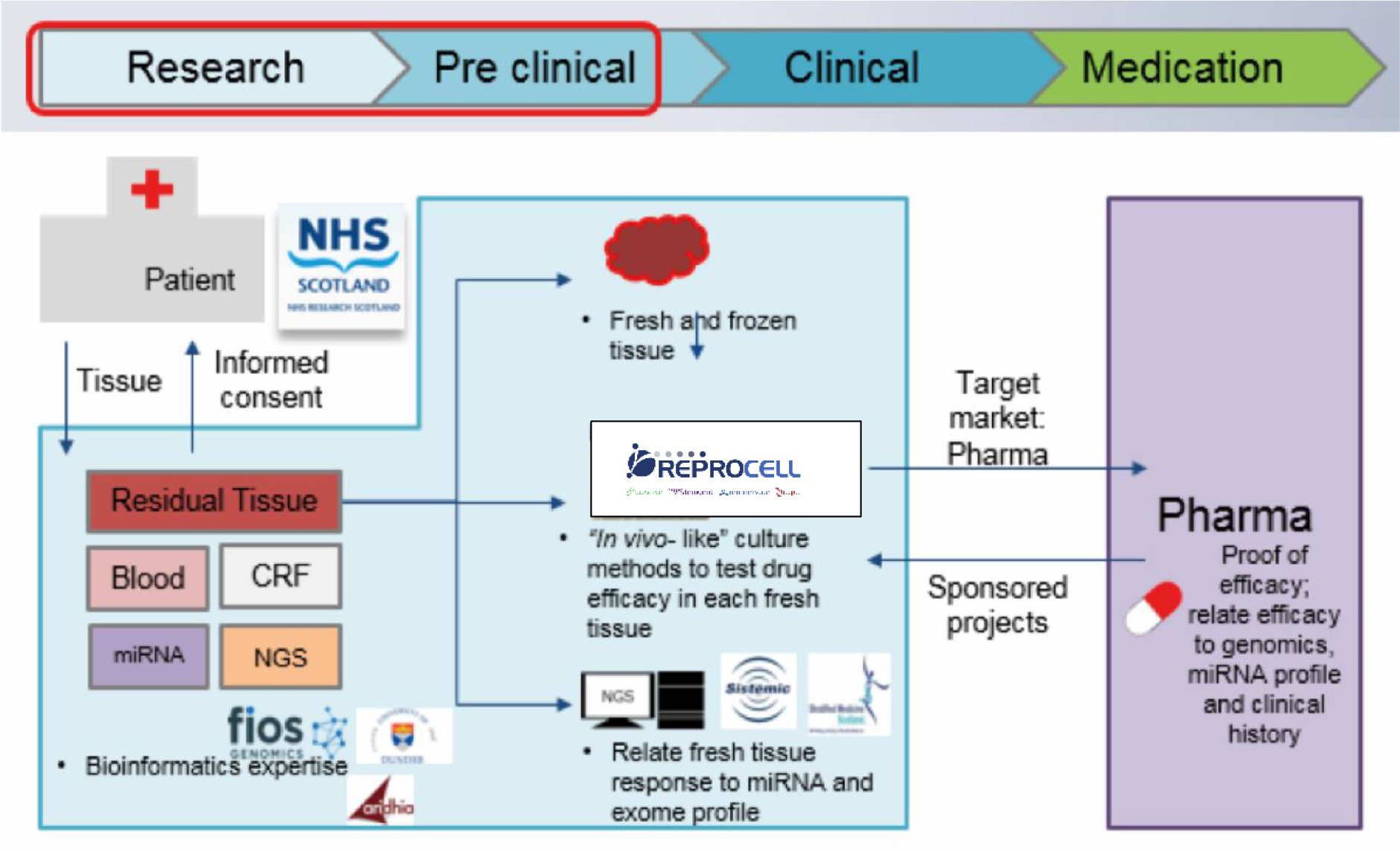
Diagram describing the precision medicine ecosystem in Scotland

**Figure 2:**
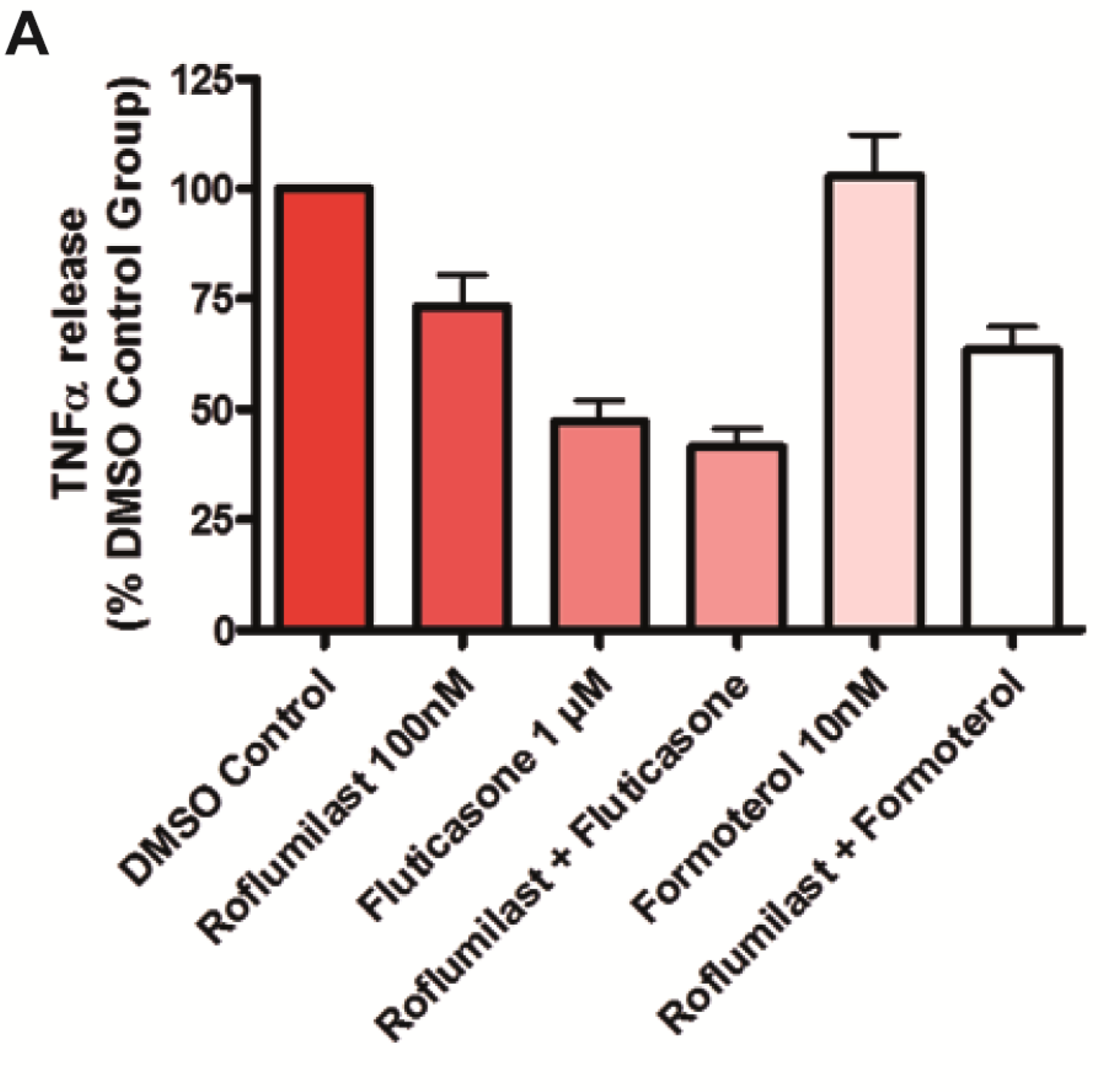
Graphs showing the effects of test articles on TNFα release from stimulated human lung parenchyma biopsies. N= 25 donors, all diagnosed with COPD. For each donor, two culture well replicates, each containing two biopsies were included in each treatment group. Data is displayed as a percentage of the corresponding DMSO control group in both graphs. A: Bar graph depicting mean + SEM TNFα release. B: Scatter graph depicting individual patient (dots) and median (thick black line) TNFα release.

Fluticasone alone or in combination with rofluminast generated the greatest inhibition of TNFα release. When the effects of monotherapy and combined therapy were compared, there was no difference in the mean reduction in TNFα levels; however, combined therapy may have resulted in a bimodal drug response across the patient sample group. Patient samples were ranked according to the level of treatment response observed in the functional organoculture assay and a bimodal pattern of response was noted in biopsies treated with roflumilast plus fluticasone (Fig 4). 12 patient samples were categorised as being high responders and 13 as low responders to the roflumilast plus fluticasone treatment.

**Figure 3.**
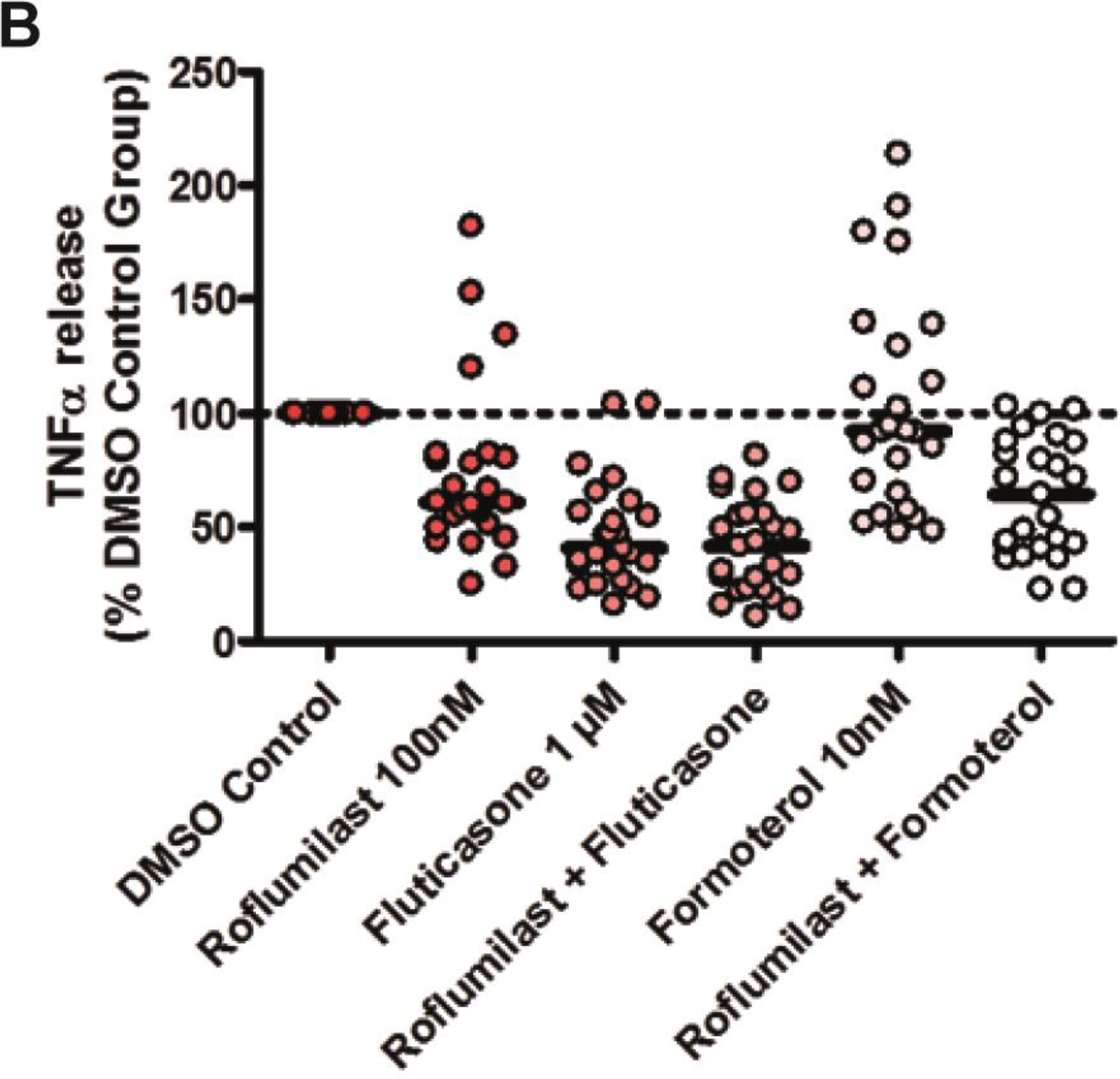
Heat map showing the results of patient demographic correlation analysis. Each parameter is assessed in relation to each other, the principal components (PC) driving variation in the data and to the organoculture assay response. Each area within the heatmap denote a p-value of association between pairs of variables from statistical tests. The statistical tests utilised depends on the property of the factors: for an association between two categorical factors, a chi-squared test was used. For an association between a categorical and a continuous factor, ANOVA was used. For an association between two continuous factors, a Spearman correlation test was used. In all cases, the resulting p-value was transformed as -log10(p) before being visualised in the confounding factors heatmap.

**Figure 4.**
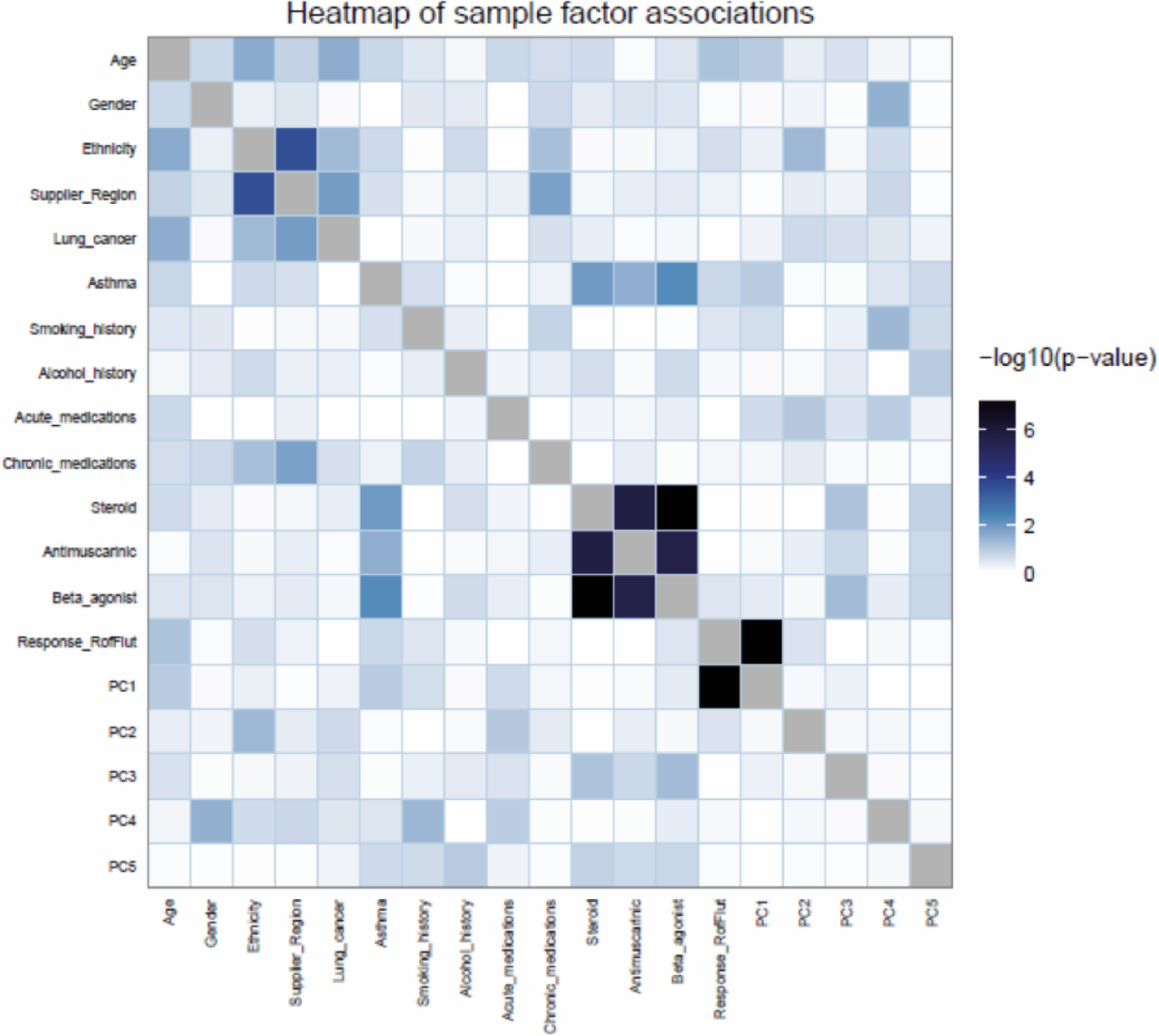
Visualisation of patient-to-patient changes in the relative levels of TNF α after combination treatment of roflumilast (100 nM) and fluticasone (1 μ M). The plot shows histogram bin counts (number of times a value falls within a given bin) as white bars as well as a smooth density in pink of the log2 ratios of TNFα release from biopsies treated with roflumilast plus fluticasone, relative to those treated with DMSO (which also includes LPS and the vehicle control), across all 25 patients. The average level in the treated biopsies is denoted by a blue dashed vertical line and the red dashed line denotes zero as this is the average level in control biopsies.

### Organoculture Bioinformatics

All patient samples passed quality control analysis as described above.

Principal component analysis was conducted to explore the relationship between the many variables.

Association analysis of patient demographic parameters and response to roflumilast plus fluticasone showed that the response was not influenced by any of the patient demographic factors such as gender or age. Treatment response was noted to be significantly associated with the first principal component, this indicates that response to roflumilast plus fluticasone is the primary trend in the data.

A strong association was observed between ethnicity and supplier region, this is however believed to be the result of one sample that was acquired from a geographical region distinct from all other regions. The ethnicity of this patient was also not replicated in any other sample.

Classes of chronic medication appear to be strongly related to each other, this is not surprising as the standard of care treatment for COPD includes combinations of the classes of drugs identified. Chronic medication appeared not to influence patient response to roflumilast plus fluticasone in the organoculture assay and is therefore not thought to be responsible for the variation in response between patients

### Exome sequence analysis

Preliminary analysis showed that all samples were of good quality, with between 38 and 57 million reads; this resulted in 36,702 to 38,065 SNPs being identified per patient sample. Merging and filtering of VCF files for high quality SNPs resulted in 101,557 SNPs being retained for the exome wide association analysis.

Hierarchical clustering and principal component analysis identified two patient samples as outliers. One of the outlying samples is described above and is thought to have resulted in a slight association with ethnicity and supplier region. There is no explanation for the second outlying sample, however as the two samples did not show any quality-related discrepancies both samples were included in downstream analysis.

Fisher’s exact test, performed in the Plink toolkit, showed that no genotypes corresponding with the identified SNPs were significantly associated with the organoculture response. However, to allow a very tentative interpretation of the results, and taking into account the low number of patients studied, an uncorrected p-value of <0.001 was chosen. With this approach a total of 30 SNPs, corresponding to 23 genes, were found to correlate with the level of TNFα release upon treatment with roflumilast plus fluticasone. A number of these genes have reported associations with COPD or other pulmonary diseases and include; HEY1^17^, SMAD3^18^, BARD1^19^ and FOXP1^20^

CYP2E1 is an inducible drug metabolising enzyme expressed in human lung tissue and has been implicated in pathological oxidative stress^21, 22^. Expression of other CYP2E1 SNPs including rs3737034 and rs2249695 were also shown to correlate with patient organoculture response. The significance level was however borderline as determined in the bioinformatic analysis (p 0.008/0.01).

Our findings suggest that genetic variation in the cytochrome p450 enzyme (CYP2E1) gene, namely SNP (rs2249695), may partly explain the observed variation in drug response. Biopsies from patients who had at least one copy of the reference allele for this SNP generally responded better to roflumilast and fluticasone co-treatment. As shown in Figure 5, mean TNFα release was inhibited by 77.6 % (homozygous reference genotype (TT)) and by 50.74 % (homozygous alternative genotype (CC)). Levels of inhibition between these two genotypes were found to be significantly different with a p value of 0.02 (unpaired, two-tailed t-test). The homozygous reference haplotype has been associated with low CYP2E1 expression^33^.

**Figure 5.**
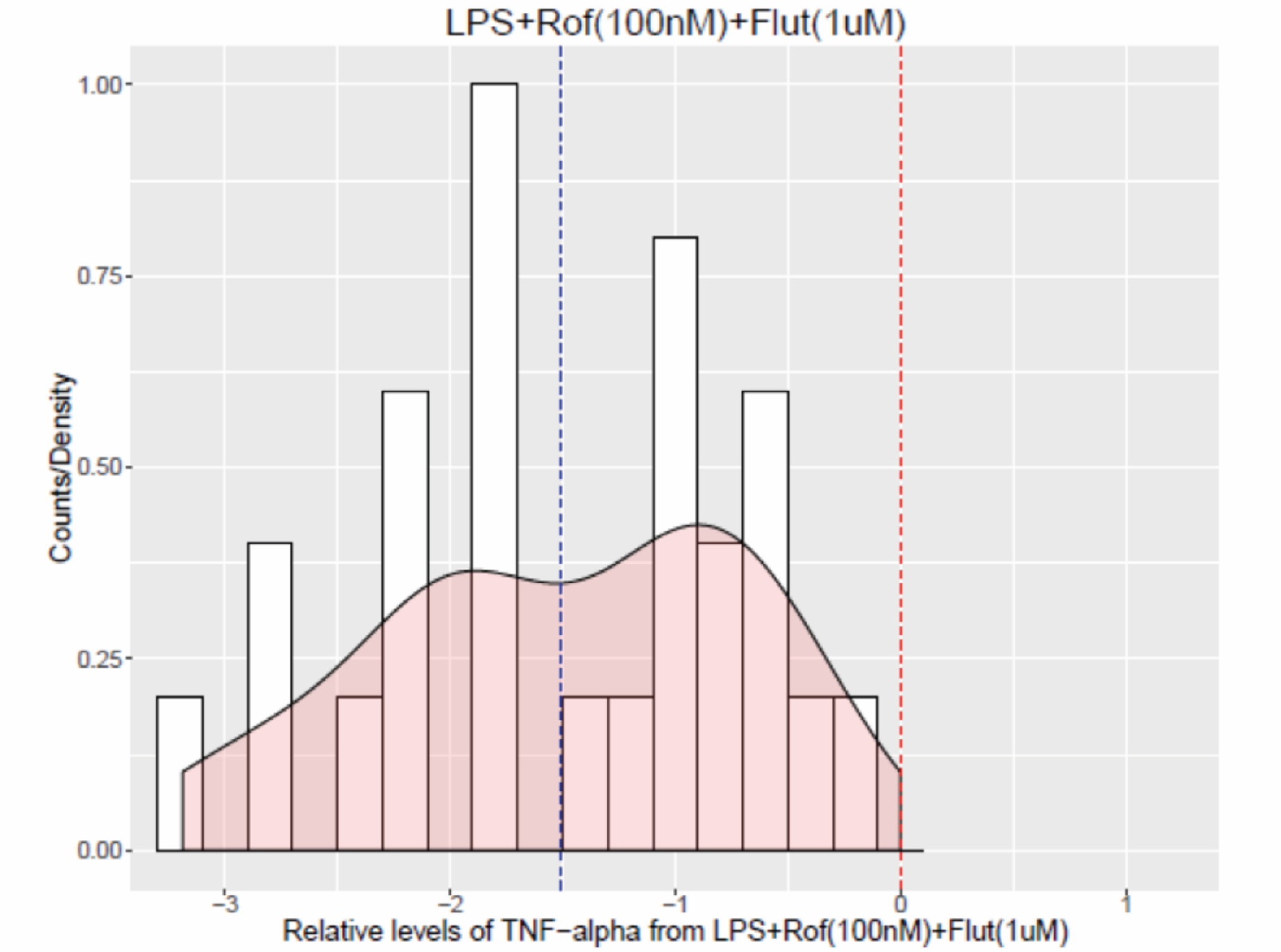
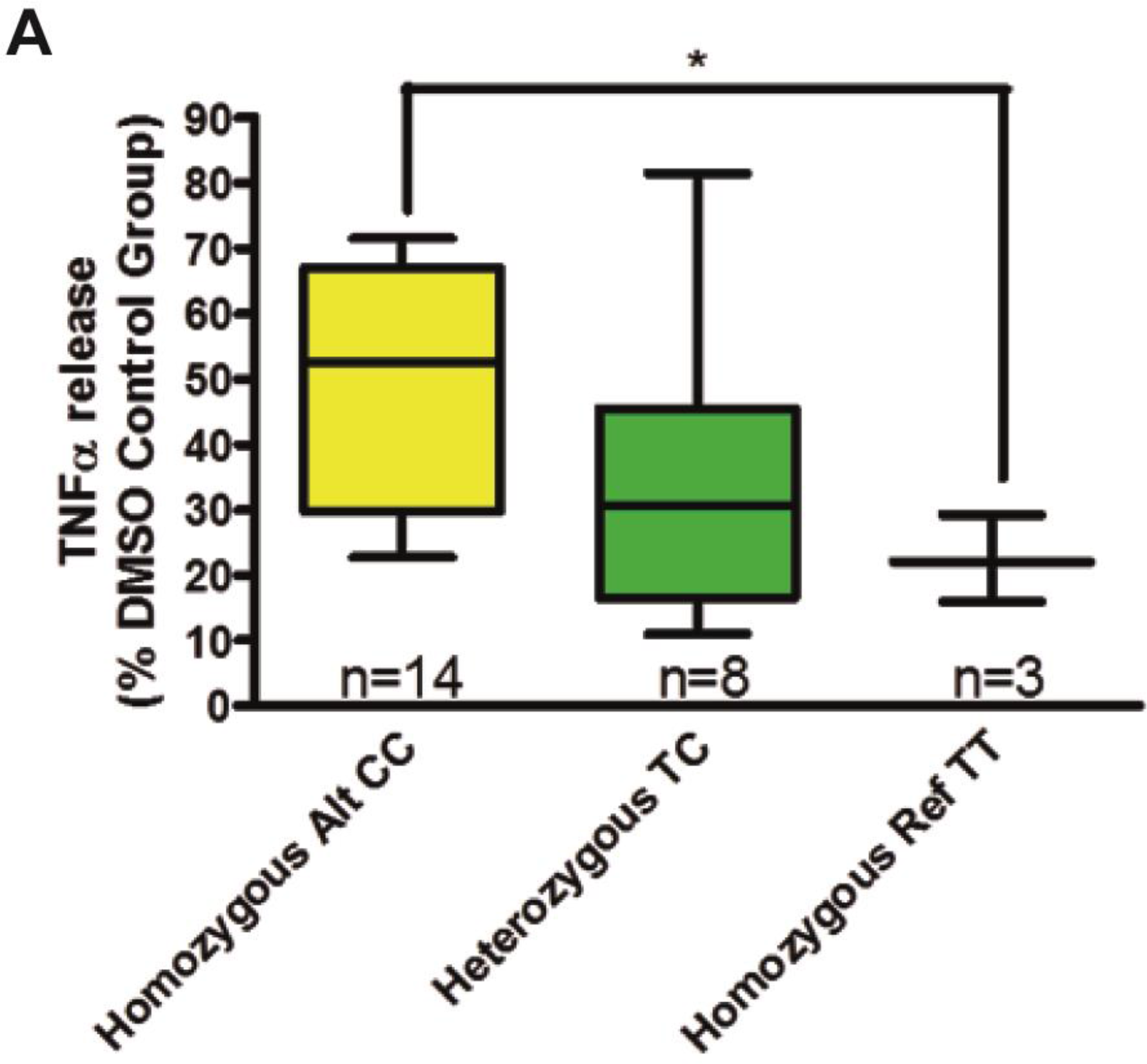
Graphs showing the relationship between CYP2E1 SNP rs2249695 genotype and TNFα release from stimulated human lung parenchyma biopsies following roflumilast and fluticasone co-treatment. Data is displayed as a percentage of the corresponding DMSO control group. Asterisks indicate significant differences (P < 0.05, for one, P < 0.01 for two and P < 0.001 for three). A: Box and whiskers graph depicting TNFα release. The 25th and 75th percentiles of each group are represented by the box with the minimum and maximum values represented by bars, the line within each box denotes the median value. B: Bar graph depicting mean + SEM TNFα release.

Genetic variation in Mothers against decapentaplegic homolog 3 (SMAD3) gene was also found to relate to patient organoculture response. As shown in Figure 6, mean TNFα release was inhibited by 66% (homozygous alternative genotype (GG)) and by 39% (heterozygous genotype). Levels of inhibition between these two genotypes were found to be significantly different with a p value of 0.0054 (unpaired, two-tailed t-test). Only two patient samples were found to have the homozygous reference haplotype (AA) and mean TNFα release was inhibited by 54 % in this group of patient samples. This level of inhibition was not significantly different to the homozygous alternative genotype or the heterozygous genotype.

**Figure 6.**
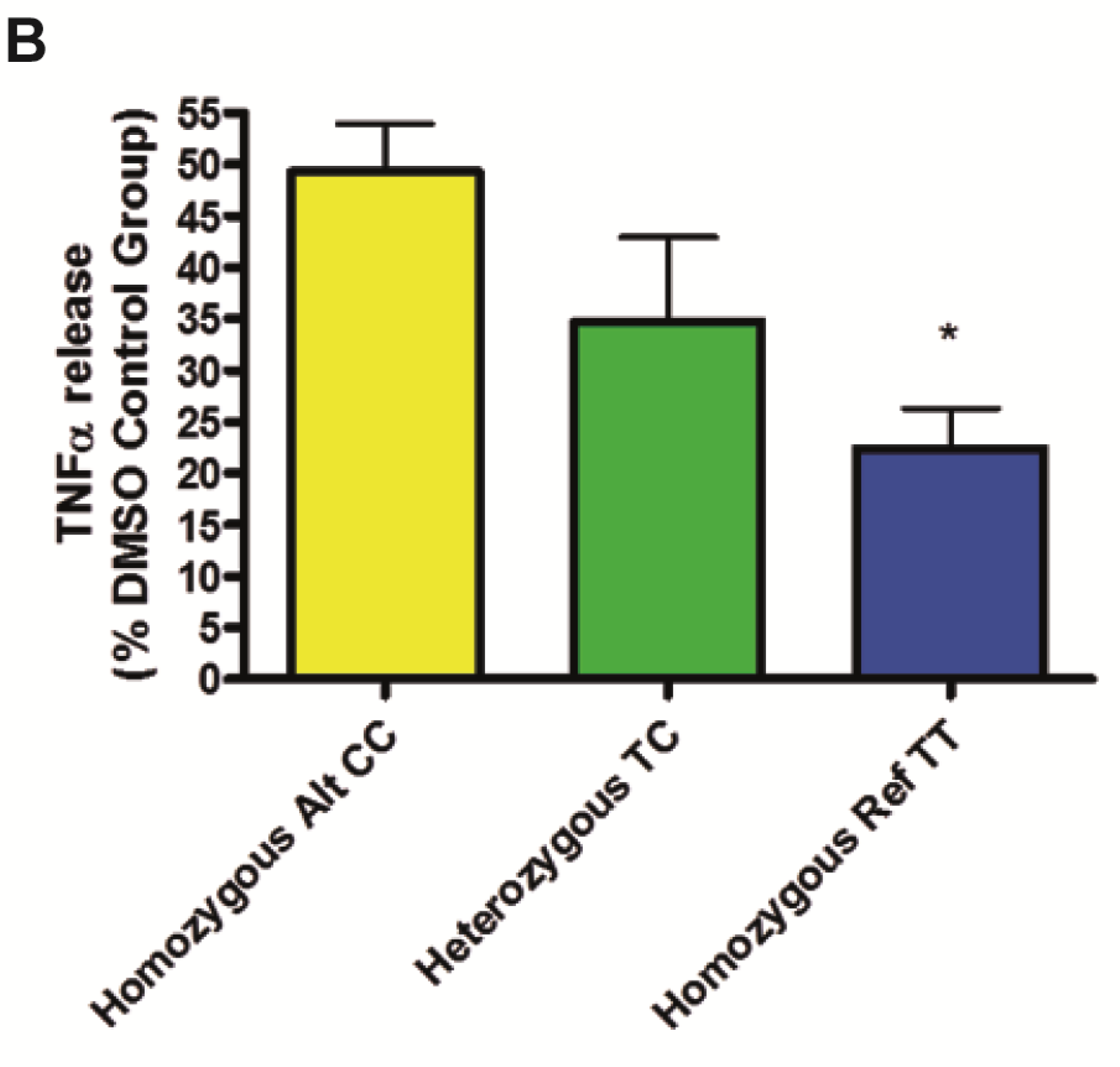
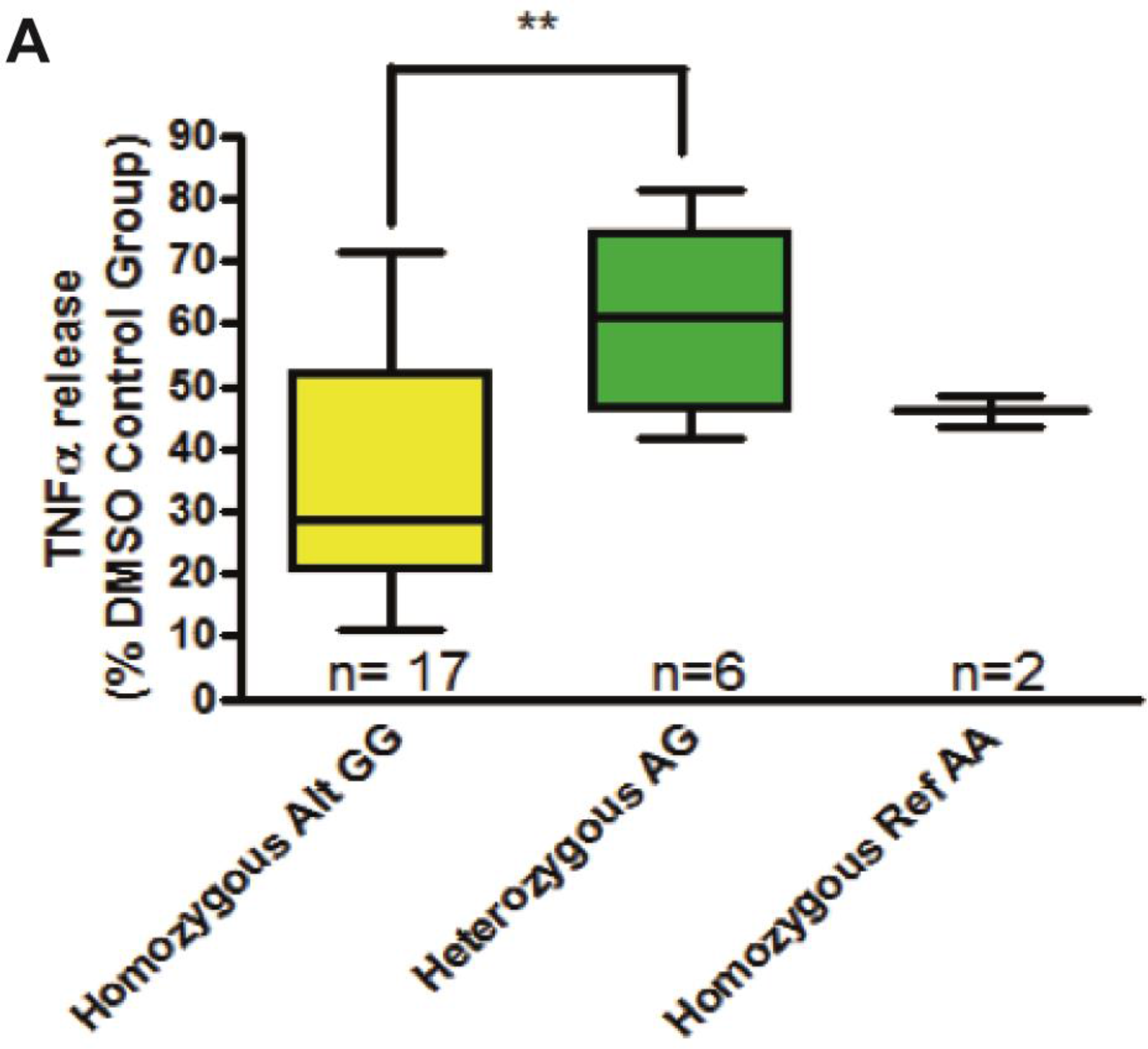
Graphs showing the relationship between SMAD3 SNP rs1065080 genotype and TNFα release from stimulated human lung parenchyma biopsies following roflumilast and fluticasone co-treatment. Data is displayed as a percentage of the corresponding DMSO control group. Asterisks indicate significant differences (P < 0.05, for one, P < 0.01 for two and P < 0.001 for three). A: Box and whiskers graph depicting TNFα release. The 25th and 75th percentiles of each group are represented by the box with the minimum and maximum values represented by bars, the line within each box denotes the median value. B: Bar graph depicting mean + SEM TNFα release.

**Figure 7.**
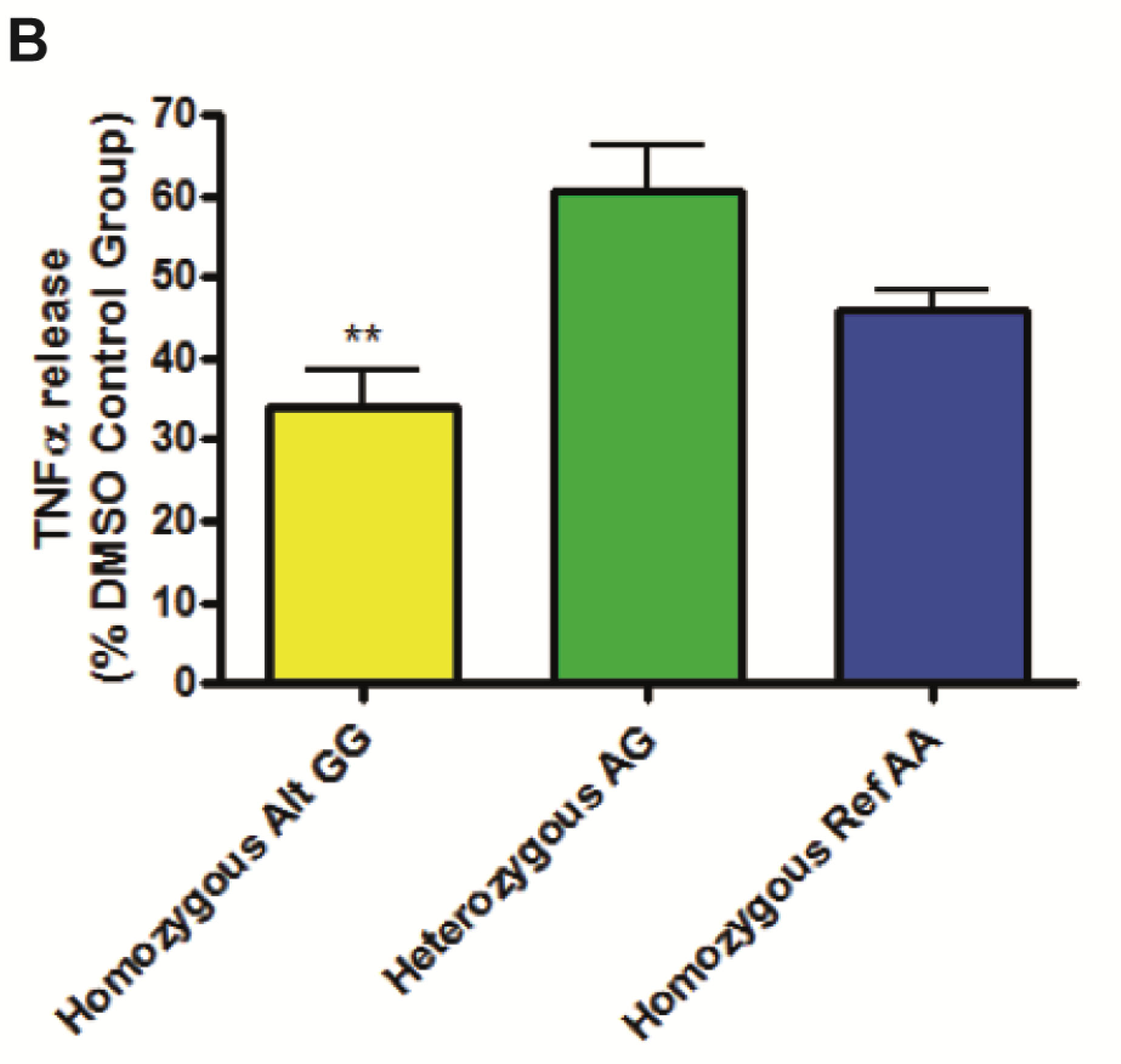
Venn diagram illustrating the overlap between genes that map to SNPs and miRNAs that are associated with patient COPD biopsy response to roflumilast plus fluticasone. Patients displayed a good response to treatment, observed by low levels of TNFα release, or a poor response as observed by high levels of TNFα release in the organoculture assay described.

The GWAS catalogue contains 624 SNPs identified in the exome sequence analysis, 4 of these SNPs are annotated in the catalogue as being associated with COPD; 6 have been associated with asthma and 4 are related to other pulmonary conditions. It was however found that no SNPs annotated in the catalogue correlated to roflumilast plus fluticasone response in this study

### miRNA analysis

RNA quality control analysis showed that isolated RNA was of high purity, 260/280 ratios ranged from 1.8 to 2.0 and RNA integrity scores ranged from 5.8 to 7.8.

All patient samples, except one, passed Agilent miRNA array quality control analysis with a rating of good to excellent. The remaining sample was flagged for evaluation and removed from subsequent bioinformatic analysis.

Statistical analysis showed that there were no specific differences in miRNA expression detected in patients that responded well in the organoculture assay compared with patients that did not respond. This analysis was performed using a p-value that had been adjusted for multiple statistical testing. For the purposes of this exemplar study, a relaxed p-value (uncorrected p <0.05) was subsequently applied. At this threshold, 181 miRNAs, mapping to 636 genes, were found to be differentially expressed in COPD patient samples that were high responders to roflumilast plus fluticasone treatment compared with samples that showed a poor response. 86 miRNAs were found to be upregulated, correlating with 47 KEGG pathways that reached statistical significance. This Enrichment analysis highlighted KEGG pathways associated with TGF-β signalling, synaptic function and fatty acid metabolism.

95 miRNAs were found to be down regulated correlating with 4 KEGG pathways that reached statistical significance. This Enrichment analysis highlighted KEGG pathways associated with long-term depression and serotonergic and GABAergic synaptic function.

1,610 GO terms were significantly associated with up-regulated miRNAs and found to be significantly enriched for pathways associated with cell ageing, specifically telomerase activity. Pathways involved in synaptic activity and T cell differentiation were also found to be upregulated.

310 GO terms were significantly associated with down-regulated miRNAs and found to be significantly enriched for pathways associated with B cell receptor activity and TGF-β production.

As discussed, bioinformatic analysis identified 30 SNPs corresponding to 23 genes (p <0.001) and 181 miRNAs (mapping to 636 genes, p <0.05) as being related to organoculture response. With further relaxation of the exome analysis p value to 0.01congruence analysis found that a total of 10 genes overlapped between the exome sequence and miRNA expression data and this overlap is higher than would be expected by chance. Overlapping genes are NTN4, IGF1R, SMAD3, EGFR, MCL1, FBN1, FGA, APP, MYO10 and IRAK3. Six overlapping genes were subject to upregulation (SMAD3, EGFR, MCL1, FBN1, FGA & APP) however the remaining 4 overlapping genes did not agree with respect to overlap direction. Absolute minor allele frequencies from the exome sequence analysis was used as a surrogate for fold changes in the SNP data. No strong correlations were found between absolute minor allele frequencies and miRNA log fold-changes. KEGG and GO enrichment analysis of the overlapping genes did not identify any common pathways or processes.

## Discussion

This study aimed to demonstrate the potential of research that combines pre-clinical functional characterisation of drug efficacy and inter-patient variation in drug responses, with state-of-the-art genomics and bioinformatics, as a new way to model precision medicine strategies at the early stages of drug development.

COPD is a highly complex condition with many clinical phenotypes. As an exemplar project, the number of patients was relatively low and findings are therefore tentative, however, this study was also designed to explore the potential for such projects during non-clinical drug development, where budgets are limited and projects exploring hundreds of patients may be too costly.

Nonetheless, clear variations in drug effectiveness were observed between patients and our preliminary experimental findings suggest that genetic polymorphisms in COPD patients may be linked to variation in response to the combination anti-inflammatory treatment, roflumilast plus fluticasone. A haplotype associated with low CYP2E1 expression was detected within the cohort of samples that responded well to treatment. It is possible that CYP2E1 expression influences response to treatment.

CYP2E1 induces production of reactive oxygen species^21, 23^ that may in turn inhibit reductions of TNFα release by various treatments. All 3 patients in the homozygous reference haplotype group were high responders to roflumilast plus fluticasone, 5 of 8 patients in the heterozygous reference haplotype group were high responders whereas 10 of 14 patients in the homozygous alternative haplotype group were low responders (Fig 5).

TGF-β and the SMAD signalling pathway have been implicated in the pathology of COPD^24–26^ and lung adenocarcinoma^27, 28^. Our results show that genetic variation in the SMAD gene (rs1065080) may influence response to fluticasone plus roflumilast. Patients that were deemed to be high responders to roflumilast plus fluticasone exclusively displayed the homozygous alternative genotype (GG), whereas only 5 of 13 patients in the poor response group displayed this genotype.

Roflumilast has been reported to inhibit TGF-β driven increases in reactive oxygen species and phosphorylation of SMAD3 by inhibiting TGF-β release^24^. If the genetic variation in SMAD3 and miRNA expression profile reported in this study alters the functioning of the pathway then this may help to explain variation in the observed organoculture response. It was however noted that no common KEGG or GO pathways were found in the bioinformatic congruence analysis.

The AnalytiXagility platform used by partners to share and interrogate the data could become a powerful resource to both academic researchers and the pharmaceutical industry.

Aridhia’s digital research platform has the potential to link the data generated in this study with available tissue, DNA and RNA for further research. A platform of this design also offers the capacity to add patients, analyses and clinical information in real time, thereby tracking patient outcome and allowing continual remodelling of the data in a secure, version controlled manner.

With ethical approval, it could be possible for researchers in the pharmaceutical industry to mine for genetic signatures or other parameters within a target disease area, for the purposes of patient selection and clinical trial support or for identifying the most appropriate pre-clinical model.

The authors acknowledge that while a very high volume of functional and genomics data was generated, the total number of patients was low for a genomics study; therefore, the scientific conclusions remain tentative but serve to demonstrate well the potential to explore patient stratification strategies at a much earlier stage by combining fresh tissue pharmacology, clinical metadata and genomics.

---

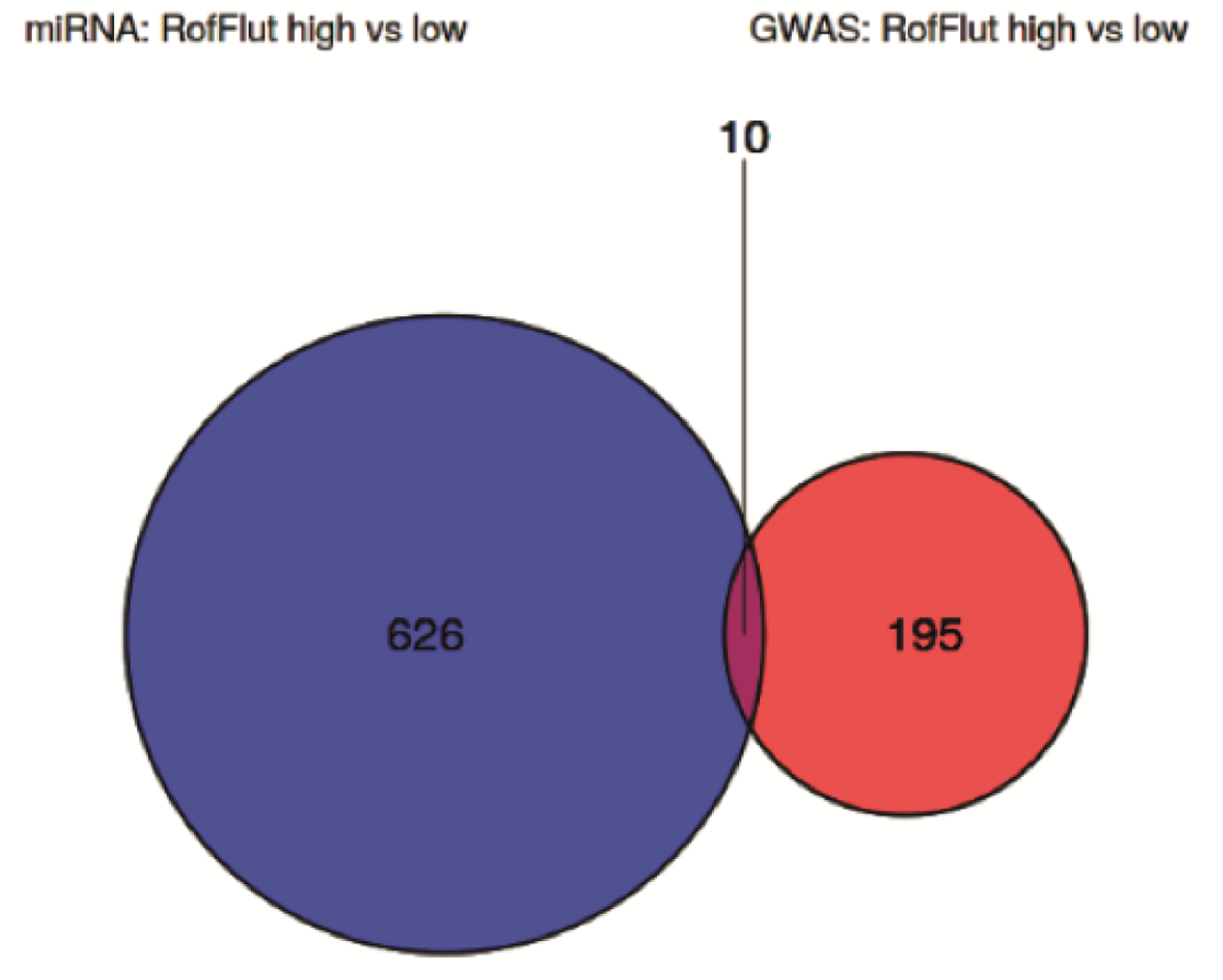

